# Amniotic Fluid Organoids As Personalized Tools For Real-Time Modeling Of The Developing Fetus

**DOI:** 10.1101/2023.10.05.561078

**Authors:** Olga Babosova, Boaz Weisz, Grace Rabinowitz, Hagai Avnet, Hagit Shani, Anat Schwartz, Linoy Batsry, Noam Pardo, Tal Elkan, David Stockheim, Tammir Jubany, Denise D. Frank, Iris Barshack, Zohar A. Dotan, Rena Levin-Klein, Pazit Beckerman, Oren Pleniceanu

**Affiliations:** Kidney Research Lab, Institute of Nephrology and Hypertension, Sheba Medical Center, Israel; Faculty of Medicine, Tel-Aviv University, Israel; Department of Obstetrics and Gynecology, Sheba Medical Center, Israel; Department of Pathology, Sheba Medical Center, Israel; Department of Urology, Sheba Medical Center, Israel; Institute of Nephrology and Hypertension, Sheba Medical Center, Israel; Sheba Medical Center Physician-Scientist Program, Israel

**Author notes:** The authors contributed equally (Olga Babosova, Boaz Weisz). These senior authors contributed equally (Rena Levin-Klein, Oren Pleniceanu).

## Abstract

Despite biomedical advances, major knowledge gaps regarding human development remain, and many developmental disorders lack effective treatment, representing a huge clinical burden. This results from fetuses being largely inaccessible for analysis. Here, we employ fetal cells in human amniotic fluid (AF) to establish personalized fetal kidney and lung organoids (AFKO and AFLO, respectively), recapitulating fetal organs at single-cell resolution. AFKO harbor key fetal kidney cell populations, including nephrogenic, urothelial and stromal, endocytose albumin, and model *PAX2*-related anomalies. Strikingly, upon injection into the nephrogenic cortex of human fetal kidney explants, AFKO-derived progenitors integrate into the host progenitor niche and contribute to developing nephrons. AFLO comprise alveolar cells and most airway cell types in a typical pseudostratified structure, upregulate surfactant expression upon corticosteroid treatment, and show functional CFTR channels. Overall, this platform represents a new personalized tool that can be applied to virtually any fetus in real-time, affording unprecedented options in studying development, uncovering mechanisms of *in utero* pathologies (e.g., congenital anomalies, infections or chemical teratogens) deciphering the developmental origins of chronic diseases, and tailoring treatments for these pathologies, as well as for prematurity-related complications. Importantly, since AF contains cells from additional tissues (e.g., skin and gastrointestinal tract), and is derived in a procedure already performed in many patients, this platform may well become a broadly applicable tool in fetal medicine.

## INTRODUCTION

Despite recent biotechnological advances, there are still considerable gaps in our understanding of human development. Likewise, the mechanisms underlying many human developmental problems, including congenital anomalies and prematurity-related complications, remain elusive^1,2^. Consequently, these disorders often lack effective treatments, accounting for 80% of non-infectious infant deaths in developed countries^3^. Moreover, although accumulating epidemiological evidence suggests a link between intrauterine development and various pathologies later in life^4^, the mechanistic basis for these associations is poorly understood.

These gaps mostly stem from the fact that human fetal tissues are largely inaccessible, making it difficult to dissect developmental processes in a specific fetus. Indeed, much of what is known on human organogenesis derives from animal studies, which suffer from interspecies differences^5^. Concomitantly, *in vitro* models generated from human fetal cells have been previously used to study development^6–9^. However, this strategy is severely limited by ethical constraints and by reliance on randomly available aborted tissue. This hinders analysis of developmental problems that are rare or do not lead to *in utero* demise, while also greatly limiting the number of available samples. Likewise, single cell-based analyses of human fetuses have greatly contributed to developmental biology^10–17^, yet suffer from the same limitations and also do not allow dynamic modeling (e.g., drug screening). Lastly, personalized fetal-like tissues can be established from pluripotent stem cells (PSCs)^18–33^. However, differentiation protocols remain suboptimal, resulting in significant off-target differentiation^34–37^. Furthermore, in ∼80% of congenital anomalies a genetic defect cannot be identified^1^, pointing to environmentally-induced epigenetic alterations as the likely cause. Since pluripotent reprogramming results in wide-spread epigenetic changes compared to the cell of origin^38–40^, PSC-derived cells may not reliably model such pathologies. Thus, novel models of human development are needed.

Here, we establish personalized tissue-specific organoids from fetal cells, by culturing human amniotic fluid (AF) cells in defined conditions. We generate amniotic fluid-derived kidney and lung organoids (AFKO and AFLO, respectively), recapitulating the fetal organ at the cellular, transcriptional and functional levels. Using congenital anomalies of the kidney and urinary tract (CAKUT) and neonatal respiratory distress syndrome (RDS) as proofs-of-concept, we show their potential in modeling and identifying therapies for developmental problems. Overall, AF organoids not only facilitate the study of normal development but also represent a novel platform enabling personalized modeling of multiple organs of virtually any fetus in real-time (i.e., while the fetus is still *in utero*). This, in turn, may allow timely implementation of precision medicine in the diagnosis and treatment of various pathologies originating in the fetal period, which until now has not been possible.

## RESULTS

### Kidney organoids can be established from AF

By the second trimester, the fetus urinates 100 ml/day^41^. Thus, we reasoned that AF cells could be used to generate organoids recapitulating fetal kidneys. Kidney development entails reciprocal interactions of the metanephric mesenchyme (MM) and ureteric bud (UB), resulting in differentiation of MM-derived nephron progenitor cells (NPC) into different nephron segments, while the UB forms the collecting system, including collecting ducts (CD) and urothelium^42^. In addition, a stromal population plays a key role in this process^43,44^.

To recapitulate this process, we collected fresh AF cells from 17–20-week pregnancies, and seeded them in basement membrane extract (BME) in a chemically defined medium (‘kidney medium’) favoring the renal lineage (**Fig. 1A**). Within 2 weeks, 3D organoids formed and gradually increased in size, displaying convoluted or spherical morphologies (**Fig. 1A-C**). The renal origin of the organoids was verified based on expression of kidney-specific factors, including several key developmental genes, such as PAX2, PAX8, CDH16, and *LHX1* (**Fig. 1A-C**)^17,45,46^. Most AFKO were epithelial, as evident by EpCAM expression, with convoluted organoids displaying complex internal structures, harboring numerous well-developed polarized tubules (**Fig. 1B**). Importantly, AFKO were viable and maintained their phenotype following cryopreservation **(Sup. Fig. S1)**. Taken together, these results demonstrate that kidney organoids can be generated from AF.

**Figure 1:**
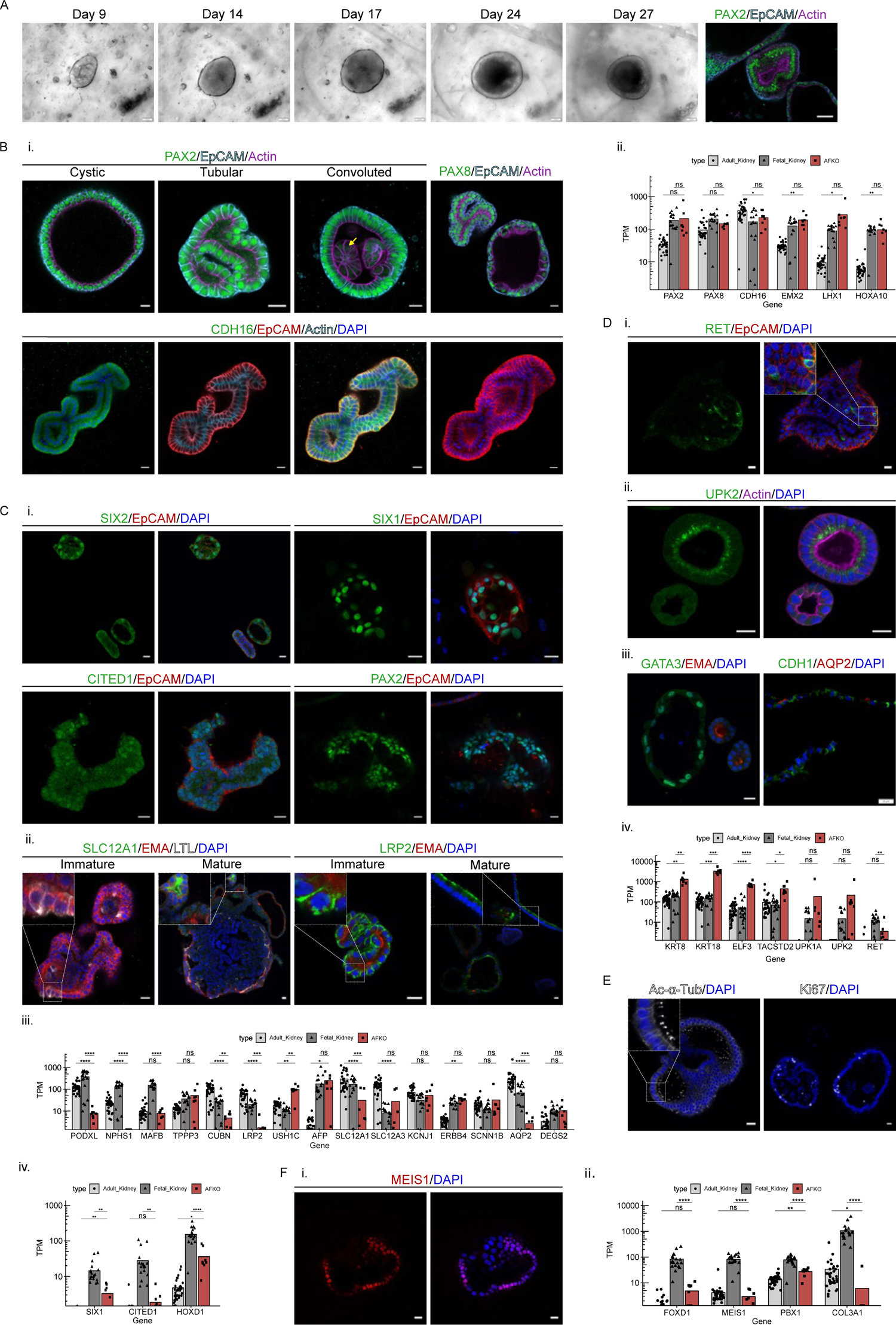
Amniotic fluid kidney organoids (AFKO) consist of cells from multiple renal compartments: **(A)** Morphology of representative AFKO. The presented organoid is PAX2^+^, confirming its kidney origin. Scale bars: 50 μm. **(B)** Expression of kidney-specific markers in AFKO. **(i.)** AFKO express PAX2, PAX8 and CDH16. PAX2 expression is shown in spherical, tubular and convoluted organoid forms; weaker PAX2 expression is seen in the inner layers of convoluted organoids (arrow). Organoid polarization is evident by the basolateral EpCAM and apical F-Actin staining. A 3D projection of EpCAM staining highlights the tubular structure of the tubular type of organoid. **(ii.)** Bulk RNA-sequencing shows expression of kidney-specific genes. **(C)** AFKO express Metanephric Mesenchyme (MM)-specific markers. **(i.)** AFKO expresses SIX2, SIX1, CITED1 at early passages (P0 - P2), while also harboring multiple PAX2^+^EpCAM^-^ cells. **(ii.)** AFKO harbor tubular structures of multiple nephron segments, as evident by expression of LTL and LRP2 (proximal tubule), SLC12A1 (loop of Henle) and EMA (distal tubule). Of note, both immature and mature tubules are seen, as evident by intracellular and membranal, as opposed to apical, marker localization, respectively. **(iii.)** Bulk RNA-seq shows expression of segment-specific genes. **(iv.)** Bulk sequencing demonstrates the expression of MM progenitor-specific genes. **(D)** AFKO express UB-specific markers, including (**i.**) UB tip marker RET, (**ii.**) urothelial marker UPK2, (**iii.**) collecting duct marker AQP2 and GATA3. **(iv.)** Bulk RNA-sequencing shows the expression of UB specific genes. **(E)** AFKO possess well-developed cilia and are proliferative, as evident by expression of apical Acetylated-α-Tubulin (cilia) and Ki67^+^ (proliferating cells). **(F)** Stromal AFKO. **(i.)** AFKO expressing the stromal marker MEIS1. **(ii.)** Bulk RNA-sequencing shows the expression of stroma specific genes. All RNA-sequencing graphs are based on 8 AFKO. Expression levels are compared to existing datasets of adult and fetal kidneys. Bar represents mean TPM (transcript per million), individual points represent individual sample TPM levels. All scale bars indicate 20 μm unless stated otherwise. Statistical significance tested by t-test. ns = not significant, *p<0.05, **p<0.01, ***p<0.001, ****p<0.0001

### AFKO harbor key fetal kidney cell lineages

Subsequently, we asked whether AFKO recapitulate the developing kidney at the cellular level, focusing first on the MM lineage. Passage (P) 0 - P2 AFKO harbored NPC expressing the master NPC transcription factor SIX1^47^, with rare cells expressing SIX2 and CITED1^48–51^, most often seen at P0 (**Fig. 1C**). SIX2 and CITED1 were undetectable at later passages, while SIX1 showed lower expression, indicating progressive differentiation. Accordingly, we detected a differentiation gradient within AFKO, ranging from a PAX2^+^EpCAM^-^ phenotype, representing undifferentiated cell types, to PAX2^+^EpCAM^+^ cells, representing tubular epithelia (**Fig. 1C**). The organoids harbored multiple nephron segments, including proximal tubule (PT, expressing LTL and LRP2), loop of Henle (LOH, expressing SLC12A1) and distal tubule (DT, expressing EMA and CDH1), often within the same organoid, and in some cases exhibited *in vivo*-like sequential presence of these segments, indicating multipotent NPC differentiation (**Fig. 1C**) As in the developing kidney^52^, we detected both mature and immature tubules, the latter lacking epithelial polarization and co-expressing several segment-specific markers. This was prominent in PTs, evident by strong early PT markers (e.g., *AFP*) alongside lower levels of maturation markers (e.g, *LRP2*) (**Fig. 1C**). We also noted expression of markers of the podocyte lineage, *MAFB* and *TPPP3*, although some mature podocytes markers (e.g., *NPHS1)* were lowly expressed, indicating presence of podocyte progenitors.

Next, we stained AFKO for RET, a marker of UB tip cells, representing UB progenitors^47^. We detected a small number of RET^+^ cells in low passage organoids (**Fig. 1D**). Accordingly, multiple organoids expressed UB markers UPK2 and GATA3, supporting a UB identity^53–55^. The expression of the CD marker AQP2, albeit in few cells, indicated a broad differentiation potential into both CD and urothelium lineages. This was supported by expression of genes representing all major UB derivatives, including cortical and medullary CD (e.g., *SCNN1B* and *TACSTD2*) and urothelium (e.g., *UPK2*)^53^. Furthermore, AFKO exhibited well-developed cilia and most harbored a small fraction of Ki67^+^ cells, indicating active proliferation (**Fig. 1E**). Importantly, we also detected stromal organoids, expressing MEIS1, a classical stromal marker^56^, which was supported by the expression of other stromal markers, such as *DCN* and *PBX1*^43,57^ and stromal progenitor marker *FOXD1*^44^ (**Fig. 1F**). Altogether, these results show that AFKO harbor the key fetal kidney lineages and sub-lineages.

### AFKO capture the cellular diversity of the developing kidney at single cell resolution

Next, we carried out single-cell RNA-sequencing (scRNA-seq) of a total of 9,264 organoid cells from two donors. Unsupervised clustering of the integrated dataset yielded 11 clusters (**Fig. 2A**). First, we validated the tissue of origin of each cluster, leveraging a dataset generated from human fetuses^17^, focusing on tissues potentially represented in AF (**Fig. 2B**). For classification, we used the top 50 markers of each organ (showing >15-fold higher expression in a given cell type versus all others; **Sup. Table ST1**). Interestingly, all clusters except for cluster 8 demonstrated renal identity, supported by the expression of renal-specific genes, such as *PAX2, PAX8* (**Fig. 2C**), and *HOXA10*. In contrast, cluster 8 (∼3.8% of cells; **Sup. Fig. S2**) had a lung-specific transcriptional signature, expressing for instance *NKX2-1* and *SFTPC* (**Fig. 2C**)^11^. Hence, we focused on kidney cells, reclustering to yield 13 distinct clusters (**Fig. 2D**), and annotating them based on marker genes identified in this and previous datasets^10,35,58,59^. This revealed 6 MM lineage clusters (0, 4, 5, 7,11, and 12), 5 UB lineage clusters (1, 2, 3, 6, and 9), 1 stromal cluster (10), and 1 proliferating cluster (8) (**Fig. 2D-F**).

**Figure 2:**
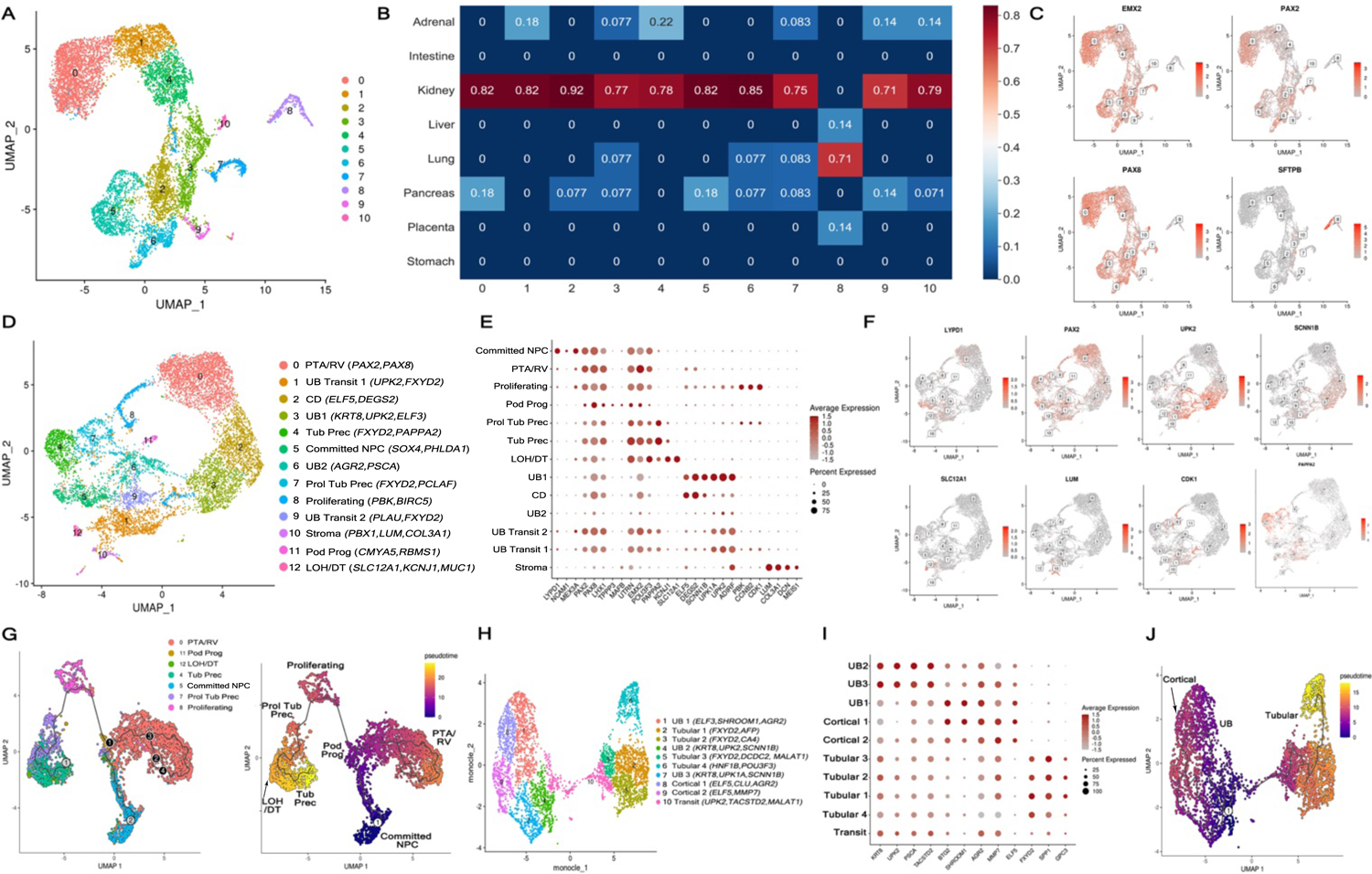
Single cell RNA-sequencing analysis AFKO: **(A)** UMAP representation AFKO cells discloses 11 clusters. **(B)** Analysis of tissue identity of each of the clusters shows all clusters to be of kidney origin, except for cluster 8, which corresponds to lung. **(C)** Expression of kidney specific (*EMX2, PAX2, PAX8*) and lung specific (*SFTPB)* genes confirms the lung identity of cluster 8 and kidney identity of all other clusters. **(D)** UMAP representation of the same dataset following removal of cluster 8, uncovers 13 cell clusters. In brackets beside every cluster are representative marker genes of the respective cluster. **(E)** Dot plot of representative marker gene expression in the different clusters. **(F)** Expression of selected marker genes. **(G)** UMAP representation of MM-derived clusters and NPC differentiation trajectory inferred by Monocle 3, with cells colored by pseudotime. **(H-J)** Analysis of UB-derived clusters. **(H)** UMAP representation of UB clusters following subclustering discloses 10 clusters. In brackets beside every cluster are representative marker genes of the respective cluster. **(I)** Dot plot of representative marker gene expression in the different clusters. **(J)** Differentiation trajectories of UB-derived clusters inferred by Monocle 3, with cells colored by pseudotime.

Among MM clusters, cluster 5 corresponded to NPC, evident by expression of *PAX2, LYPD1,* and *VIM.* This cluster did not express early NPC markers (e.g., *SIX2),* as these were P1 AFKO, thus termed ‘committed NPC’. Notably, *NCAM1*, previously reported to mark NPC in human fetal kidneys^58,60,61^ was identified as a marker gene of this cluster. Cluster 0 was identified as pre-tubular aggregate/renal vesicle (PTA/RV) stage, based on *PAX8* and *LHX1* expression, representing NPC descendants that still harbor wide differentiation potential. Clusters 4 and 7 corresponded to ‘tubular precursors’^10^, expressing markers of both PT and DT, such as *PAPPA2* and *KCNJ1*. Cluster 7 also expressed high levels of proliferation genes and was named ‘proliferating tubular precursors’. Cluster 12 exhibited a more differentiated phenotype, consistent with a common precursor of LOH and DT (‘LOH/DT’) based on expression of markers of both segments (e.g., *SLC12A1* and *MUC1*). Lastly, cluster 11 expressed markers of the podocyte lineage (e.g., *MAFB*), although markers of mature podocytes (e.g., *NPHS2*) were absent, confirming a podocyte progenitor identity. The proliferating cluster (8) exhibited strong *PAX2* and *LHX1* expression, suggesting MM origin.

UB clusters were identified based on expression of *KRT8, ELF3, ELF5* and *UPK2*^55^. Two clusters, ‘UB1’ and ‘UB2’, expressed markers corresponding to multiple UB-derived cell types, including cortical (e.g., *ELF5*) and medullary CD (e.g., *PSCA)* and urothelium (e.g., *UPK2* and *UPK3A*)^53^. In contrast, cluster 2 expressed predominantly CD markers. Hence, ‘UB’ clusters likely represent less differentiated UB cells, while the ‘CD’ cluster represents CD-primed UB cells. Interestingly, 2 clusters expressing UB markers also expressed MM-lineage markers, suggesting a transitory phenotype between UB and MM, termed ‘UB-transit’.

Lastly, cluster 10 comprised cells expressing renal stromal markers, such as *MEIS1* and *PBX1*. Although it did not express markers of stromal progenitors (*FOXD1* or *TBX18*)^44,62–64^ it demonstrated *OSR1* expression, indicating its renal origin. Altogether, these results show that AFKO harbor the myriad cell types of the human fetal kidney.

### AFKO recapitulate developmental trajectories of the human fetal kidney

Subsequently, we dissected differentiation trajectories in the MM lineage by analyzing cells annotated as belonging to this lineage (**see Methods**) (**Fig. 2G**). This uncovered a pseudo-time trajectory starting in NPC and continuing through the PTA/RV cluster. Subsequently, PTA/RV give rise to podocyte progenitors, or alternatively go through the proliferative state and commit to the tubular lineage, passing first through the ‘proliferative tubular precursor’ state and generate tubular precursors. Accordingly, LOH/DT cells were at the end of this trajectory.

We then performed ligand-receptor interactions to identify intercellular interactions in AFKO **(Sup. Fig. S3)**, focusing on NPC, PTA/RV, tubular precursor, UB, podocyte progenitor, and stromal clusters. Interestingly, we detected FGFR2-FGF9 interactions involving primarily NPC and PTA/RV cells, consistent with the role of FGF signaling in NPC self-renewal^65,66^. Likewise, JAG1-NOTCH2 interaction was active in these clusters, consistent with its role in NPC differentiation^67–69^. We also noted enrichment for FZD1-LRP5-WNT7B interaction in UB-related interactions, including with NPC, where UB-derived WNT7B interacted with FZD1 in NPC, in accordance with the role of UB-derived Wnt ligands in nephrogenesis^70,71^.

To better define UB cell types, we subclustered and annotated UB clusters using a recent dataset^53^, yielding 10 clusters (**Fig. 2H**). Three clusters (‘UB1’, ‘UB2’, and ‘UB3’) expressed markers encompassing all types of ureteric epithelia, including cortical (e.g., *AGR2*), outer medullary (e.g., *SHROOM1*) and inner medullary (e.g., *PSCA*; **Fig. 2I**). Thus, these clusters likely correspond to UB cells with wide differentiation potential. In contrast, clusters ‘Cortical 1’ and ‘Cortical 2’ expressed mostly cortical ureteric epithelium-related genes, including principal cell marker *ELF5*. Interestingly, four ‘tubular’ clusters showed relatively strong expression of tubular markers (e.g., *FXYD2* and *SPP1*), confirming that some UB cells acquire tubular features. Lastly, cluster 10 exhibited an intermediate signature between ‘UB’ and ‘Tubular’ clusters, likely representing a transitory state. Pseudotime analysis indicated that UB cells in AFKO give rise to cortical ureteric epithelial cells, or alternatively, acquire a tubular-like phenotype (**Fig. 2J**). Taken together, these results show that AFKO mirror key cell states and trajectories of the fetal kidney.

### AFKO exhibit tubular functionality and allow CAKUT modeling

To determine AFKO functionality, we assessed their ability to uptake albumin, a clinically-relevant function of PT cells^72^ exerted by the LRP2-Cubilin complex. Since LRP2 is lowly expressed in AFKO and mostly shows non-luminal localization (**Fig. 1C**), we first modified culture conditions **(Sup. Fig. S4A)** to induce proximal differentiation, including LRP2 upregulation^73^ (**Fig. 3A**). We then incubated AFKO with fluorescently-labeled albumin at 37°C (allowing active transport) or 4°C (active transport inhibited) (**Fig. 3B-C**). AFKO incubated at 37°C demonstrated significantly greater albumin uptake, confirming active albumin uptake.

**Figure 3:**
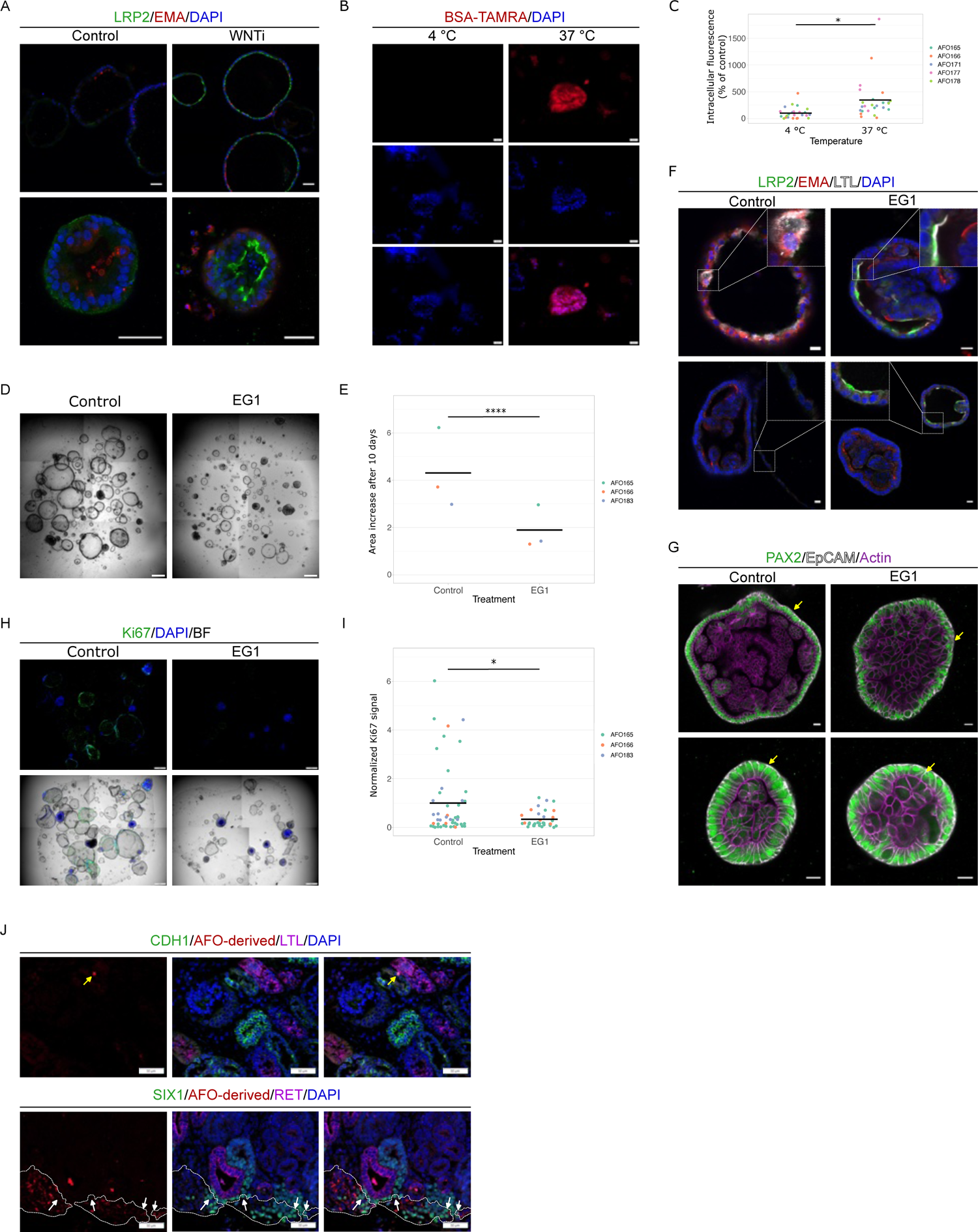
AFKO recapitulate kidney functionality and serve as a model of kidney anomalies: **(A)** WNT inhibition induces proximal differentiation of AFKO, as evident by LRP2 induction relative to DMSO-treated control following 3 days of treatment. Scale bars: 50 μm. **(B-C)** AFKO endocytose albumin as shown in the BSA-Tetramethylrhodamine (BSA-TAMRA) assay of LRP2/CUBN transport. **(B)** Representative images of intracellular BSA-TAMRA (red) after 2 hours of incubation at 37°C, demonstrating active BSA transport. Active BSA transport is inhibited at 4°C. Scale bars: 20 μm. **(C)** Quantification of intracellular BSA-TAMRA fluorescence. Normalized quantification of n = 5 independent experiments, with each experiment including n = 5 organoid area measurements. Lines indicate means per condition. *p = 0.0179 - Mann Whitney test on all measurements. **(D-I)** PAX2 inhibition in AFKO generates an anomalous phenotype. AFKO were treated with the PAX2 inhibitor EG1 and compared to DMSO-treated AFKO. **(D)** Representative images of AFKO following 10 days of treatment, showing reduced organoid size. Scale bars: 500 μm. **(E)** Quantification of AFKO area fold increase 10 days after treatment. n=3 independent experiments, with each experiment including n = 20 organoid area measurements. Lines indicate means per condition. ****p = 1.348e-07 - Mann Whitney test on all measurements. **(F)** EG1-treated AFKO show enhanced maturation, as evident by the apical localization of LTL and LRP2, as opposed to the intracellular LTL localization and lack of LRP2 in DMSO-treated controls. Scale bars: 20 μm. **(G)** EG1-treated AFKO show loss of epithelial integrity (arrows). Scale bars: 20 μm. **(H-I)** EG1-treated AFKO show reduced proliferation, as evident by Ki67 staining. **(H)** Representative image of an entire well containing Ki67-stained AFKOs. Scale bars: 500 μm. **(I)** Quantification of Ki67 fluorescent staining in EG1- and DMSO-treated organoids. Normalized quantification of n = 3 independent experiments, with a total of n ≥ 35 organoids in each condition. Lines indicate means per condition. *p = 0.0198 - Mann Whitney test on all measurements. **(J)** AFKO cells dyed with celltracker Red CMTPX injected into human fetal kidney explants show integration into tubular structures, as shown by the yellow arrow pointing to an AFKO cell in a CDH1^+^ tubule. AFKO cells also integrate into the native nephrogenic niche adjacent to RET^+^ UB (marked by a white line), with some of these cells showing NPC identity, as evident by SIX1 expression (white arrows). Scale bars: 50 μm.

We next asked whether AFKO could model CAKUT, focusing on renal-coloboma syndrome caused by PAX2 loss-of-function, and manifesting as renal hypodysplasia. Interestingly, studies using PSC-derived kidney organoids did not find a major defect in tubular epithelialization following PAX2 knockout^74^. To this end, we treated AFKO with the PAX2 inhibitor EG1^75^. Following 10 days of exposure to EG1, the organoids demonstrated reduced growth compared to mock-treated AFKO, while breast organoids were unaffected (**Fig. 3D-E**, **Sup. Fig. S4B**). Histologically, we found epithelial structure disruption and decreased Ki67 expression as in previous studies^76,77^, as well as enhanced tubular marker expression (**Fig 3F-I**), pointing to premature differentiation as a potential cause of the phenotype.

### AFKO show robust differentiation potential *in vivo*

Since they contain undifferentiated kidney cells, AFKO may afford a potential source for cell therapy. To test this option, we dissociated AFKO from two donors into single cells, labeled them with cell tracker Red CMTPX dye and injected them into human fetal kidney explants derived from healthy 2nd trimester fetuses (see methods). Following two days, explants were fixed and analyzed by immunostaining (**Fig. 3J**). Remarkably, we detected multiple SIX1^+^ NPC originating from the organoids integrated within the nephrogenic cortex of the injected kidney, adjacent to RET^+^ UB tips. In addition, a subset of injected cells integrated into epithelial structures of the injected kidney. Taken together, these results demonstrate the ability of AFKO-derived cells to integrate into the host nephrogenic niche as NPC within the native NPC population, and to contribute to developing nephrons.

### AFLO recapitulate the human developing lung

Subsequently, we modified culture conditions to enrich for lung progenitors. Using a defined ‘lung medium’, we obtained cystic organoids, expressing the canonical lung marker NKX2-1^78–80^ (**Fig. 4A-B,H**). A subset of cells co-expressed SOX2 and SOX9, indicating distal tip identity, representing fetal progenitors that initially gives rise to proximal (airway) progeny and during weeks 17-20 become gradually committed to distal (alveolar) fates^6,81,82^. Accordingly, distinct SOX2^+^ proximal and SOX9^+^ distal niches were seen within AFLO (**Fig. 4C**). Notably, cystic AFLO, composed of a simple epithelium, gradually acquired complex morphologies, reflected microscopically by typical pseudostratified epithelium, with p63^+^ basal cells residing beneath differentiated airway cells, including SCGB1A1^+^ club, FOXJ1^+^ ciliated and CALCA^+^ neuroendocrine cells (**Fig. 4D-F,H**). Another organoid type acquired a convoluted morphology, demonstrating an even more complex internal organization, harboring ‘airway-like’ structures, whose lumen contained secreted MUC1 (**Fig. 4D**). Adjacent to these structures we identified SCGB1A1^+^ club cells. Concomitantly, we detected alveolar organoids, usually expressing markers of both alveolar type 1 (AT1, HOPX) and 2 (AT2, proSP-B and proSP-C) cells, with some cells expressing both markers, indicating common alveolar progenitors (**Fig. 4G-H**), as reported in human fetal lungs^11,83^. Strikingly, some AFLO demonstrated a highly intricate structure, comprising both airway and alveolar cells (**Fig. 4J**). Transmission electron microscopy (TEM) confirmed presence of lamellar bodies, indicating AT2 identity (**Fig. 4I**).

**Figure 4:**
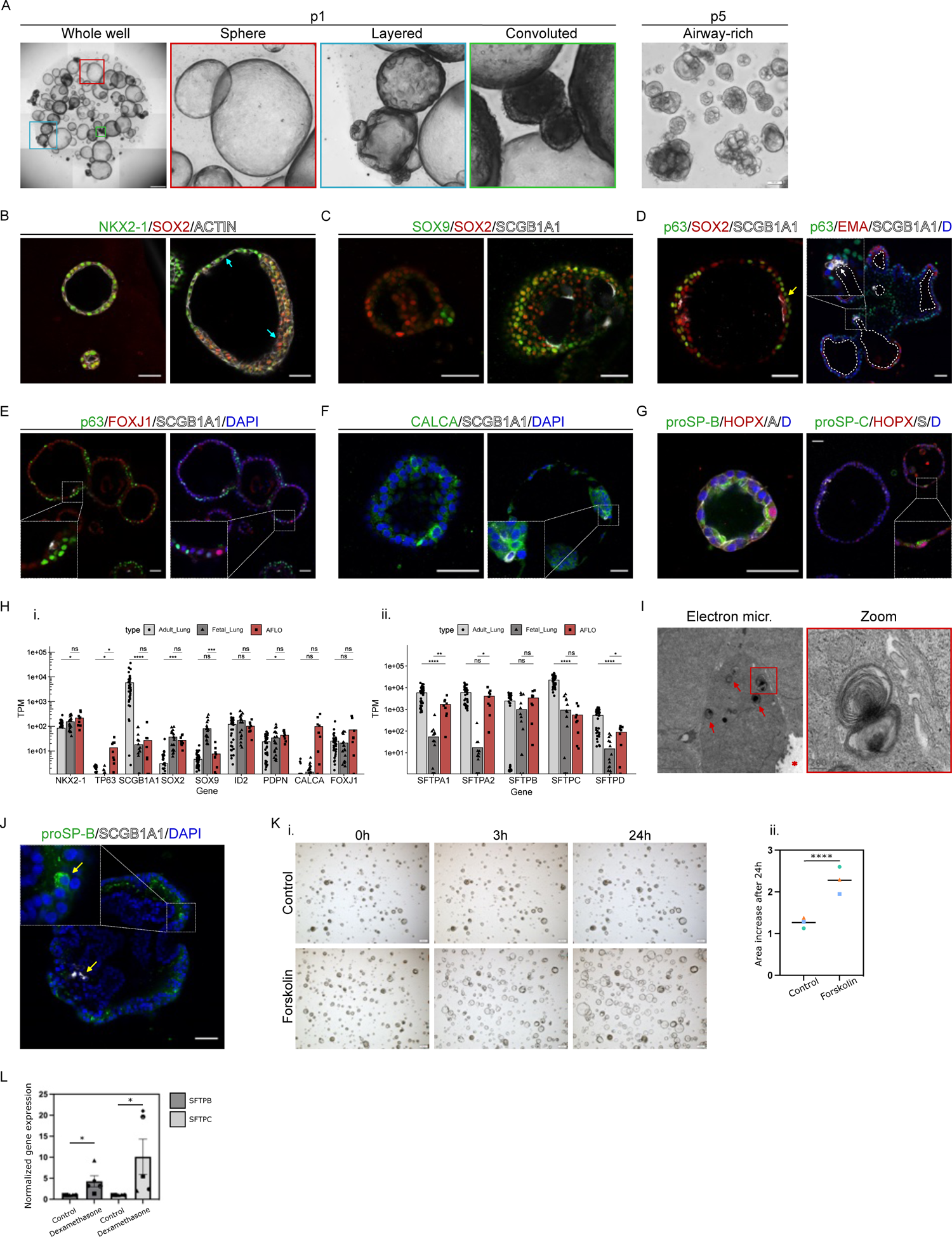
AFLO harbor proximal and distal lung lineages and show functionality *in vitro*: **(A)** Bright field images of AFLO at P1, showing different stages of organoid development. Scale bar: 1 mm. The three described stages of sphere, layered and convoluted forms are enlarged. Another form, containing airway-like structures, usually observed at later passages is shown at P5. Scale bar: 100 μm. **(B)** AFLO express the lung-specific marker NKX2-1. Once spheric AFLO become layered, SOX2 is upregulated and NKX2-1 is downregulated (cyan arrows). Scale bar: 50 μm. **(C)** AFLO harbor both SOX2^+^ proximal and SOX9^+^ distal progenitors, including double positive SOX2^+^SOX9^+^ cells. Scale bar: 50 μm. **(D)** AFLO harbor all major airway cell types and display a typical pseudostratified epithelial structure. P63^+^ basal cells co-stain with SOX2 and localize to the basal layer (yellow arrow). SCGB1A1^+^ club cells are also SOX2^+^ and assume a luminal position in airway structures (white arrow). Airway structures are positive for mucin-1 (EMA). Lumen is demarcated by dotted white lines; D, DAPI. **(E)** FOXJ1^+^ ciliated cells are seen in AFLO. **(F)** AFLO harbor CALCA^+^ neuroendocrine cells. **(G)** Alveolar AFLO express proSP-B, proSP-C and HOPX. A, Actin, S, SCGB1A1, D, DAPI. **(H)** Bulk RNA-seq of 9 AFLO. **(i.)** Lung developmental and airway lineage genes. **(ii.)** Alveolar lineage genes. Expression levels (mean TPM) are compared to existing datasets of adult and fetal lungs. ns = not significant, *p<0.05, **p<0.01, ***p<0.001, ****p<0.0001 - Student’s t-test. **(I)** Representative transmission electron microscopy (TEM) of AFLO section. Red arrows indicate lamellar bodies. Red star indicates the organoid lumen, scale bar: 2 μm. Right panel shows enlarged detail of the lamellar body marked by a red box in the left panel. Scale bar: 200 nm. **(J)** Immunostaining of airway-specific SCGB1A1 marker and alveolar-specific proSP-B marker (yellow arrows), demonstrate their expression in the same organoid structure. Scale bar: 50 μm. **(K)** AFLO swell in response to forskolin relative to DMSO-treated control. **(i.)** Bright field images of AFLO incubated for indicated times. Scale bar: 200 μm. **(ii.)** Measurement of fold area increase of DMSO- and forskolin-treated AFLO following 24 hour treatment. Means of n = 3 independent experiments, each dot represents an experiment, with each experiment n = 20 organoid area measurements. Lines indicate means per condition. ****p = 2.845e-14 - Mann Whitney test on all measurements. **(L)** Dexamethasone treatment significantly increases the expression of *SFTPB* and *SFTPC* in AFLO. Dotbar plot represents n = 5 biological replicates, each dot represents a different donor. Bars represent mean, error bars standard error of the mean. *SFTPB* *p = 0.0152, *SFTPC* *p = 0.0201 - paired two-tailed t-test. All scale bars indicate 50 μm unless stated otherwise.

We next analyzed AFLO using scRNA-seq, sequencing 2 samples containing 26,101 cells after quality control **(see Methods)**. Unsupervised clustering of the integrated dataset resulted in 12 clusters (**Fig. 5A, Sup. Fig. S5A)**. Using the same approach as in AFKO, we verified the lung origins of all clusters (**Fig. 5B**). We next characterized clusters based on their markers. We detected the presence of tip-like SOX2^+^GATA6^+^ cells^6,82–84^, alongside multiple airway cell types, including basal, club, ciliated, neuroendocrine, and secretory, identified based on expression of *TP63*, *SCGB1A1*, *FOXJ1*, *CALCA*, and *KRT6B*, respectively (**Fig. 5A**). As expected, we noted 2 alveolar clusters showing a common AT1-AT2 identity, based on co-expression of markers of both lineages (e.g., *AQP4* and *SFTA2*). Interestingly, ‘basal 3’ cluster exhibited a transitory state between basal cells (representing airway progenitors) and their differentiated progeny, based on low expression of classical basal markers (e.g., *TP63*) and high expression of more differentiated cells (e.g., *MUC5B* and *SCGB3A1*), potentially corresponding to the suprabasal/luminal progenitor cell type^85^. Trajectory analysis unraveled two main trajectories starting from tip-like cells, corresponding to airway and alveolar lineages. The former involved acquisition of a basal cell state with subsequent transition into secretory, neuroendocrine and ciliated identity. Interestingly, while some club cells were seen within this trajectory, a substantial number of them were connected to the alveolar trajectory, potentially corresponding to similar bipotent cell types recently identified in the adult human lung^86,87^. Collectively, these results show that AFLO recapitulate cell types and lineage relationships of the developing lung.

**Figure 5:**
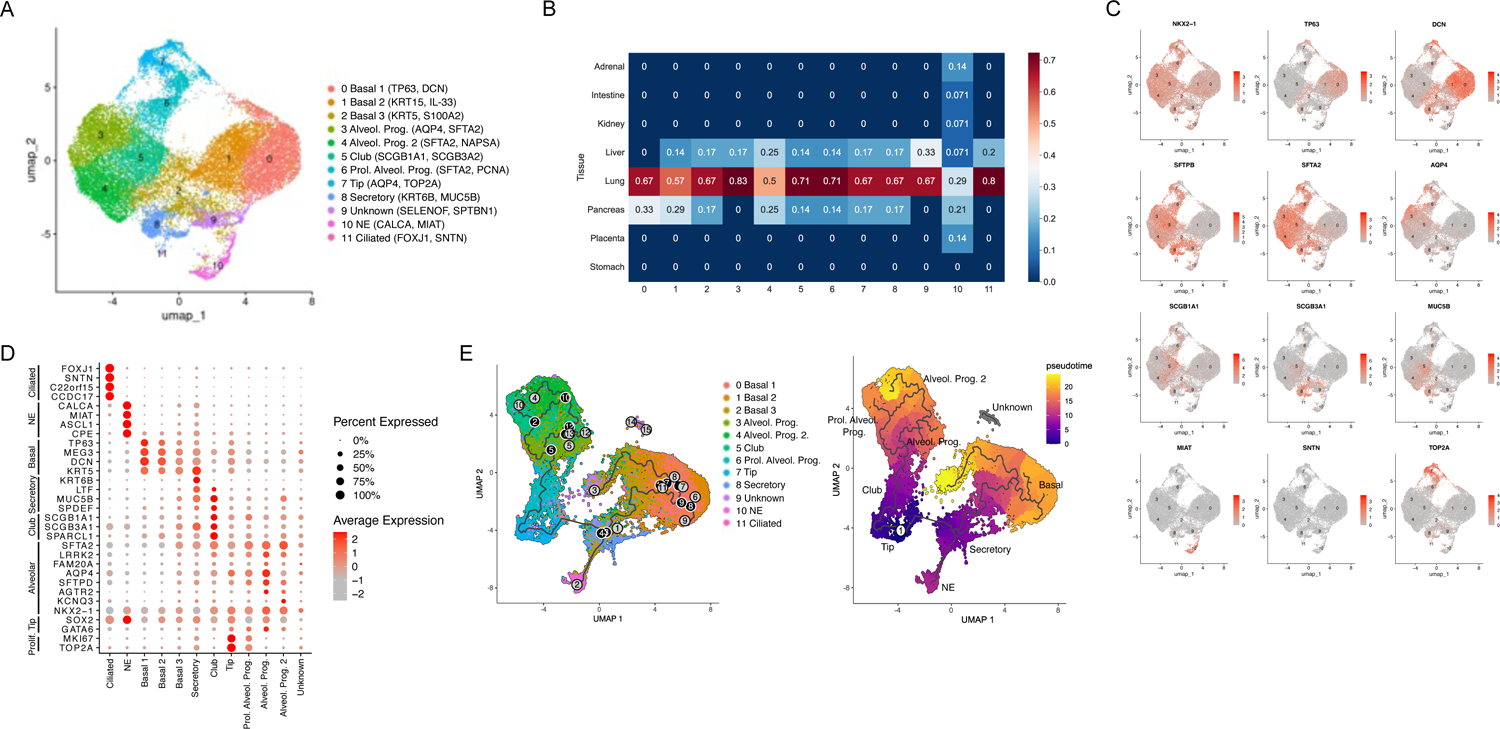
Single cell RNA-seq analysis AFLO: **(A)** UMAP representation of AFLO-derived cells discloses 11 clusters. In brackets beside every cluster are representative marker genes of the respective cluster. **(B)** Analysis of tissue identity of each of the clusters shows all clusters to be of lung origin. **(C)** Expression of selected marker genes. **(D)** Dot plot of representative marker gene expression in the different clusters. **(E)** Left panel: UMAP representation of trajectory inferred by Monocle 3. Right panel: Pseudotime starting at cluster 7 (Tip).

### AFLO are functional and can serve to model developmental problems

To determine AFLO function, we treated them with the CFTR inhibitor Forskolin. This resulted in significant swelling, indicating functional CFTR channels^88^ (**Fig. 4K**). Next, we tested their potential in modeling neonatal RDS, a common condition in preterms, which is associated with significant morbidity and mortality, and results from AT2 immaturity with consequent reduced levels of surfactant proteins. RDS is treated with prenatal corticosteroid administration^89^, which has partial success and variable patient response^90^. We thus treated AFLO for 72 hours with dexamethasone. Strikingly, we noted a significant increase in both *SFTPB* and *SFTPC*, noting variable response rates, as in patients (**Fig. 4L**). Taken together, these results indicate that AFLO are functional and facilitate the study of developmental lung problems.

### AFKO and AFLO show distinct organ-specific transcriptional signatures

To assess the robustness of our organoid platform, we carried out bulk RNA-seq using the MARS-seq method^91^ on matched AFKO and AFLO generated from 4 donors at early (P1-2) and later (P4) passages, as well as on kidney tubuloids^92^, generated from 4 adult donors, and 1 unmatched AFLO. Remarkably, unsupervised euclidean-based clustering, as well as PCA analysis, clearly showed separation based on tissue of origin, with AFKO clustering together with tubuloids, and separately from AFLO (**Fig. 6A-B**). Organoids derived from the same donor clustered based on the differentiation protocol used, superseding donor identity. Differentially expressed gene (DEG) analysis between all kidney-related organoids and AFLO revealed large and consistent differences between these samples, with 658 genes upregulated in kidney samples, and 618 in lung samples (**Fig. 6C, Sup. Table ST2**). Interestingly, while most AFKO clustered together, some were more transcriptionally similar to tubuloids, indicating that different maturation levels are obtained in different donors. Pathway analysis of DEGs between AFKO and AFLO revealed clear enrichment for organ-specific pathways (**Fig. 6D**). For instance, processes enriched in AFKO included ‘renal system development’ and ‘ureteric bud development’, while enriched cell types included ‘proximal tubules’, ‘distal tubules’, and ‘collecting ducts’, confirming that AFKO capture key cell types of the developing kidney. In contrast, AFLO showed enrichment for ‘respiratory system development’ and ‘lung alveolus development’, corroborating scRNA-seq results.

**Figure 6:**
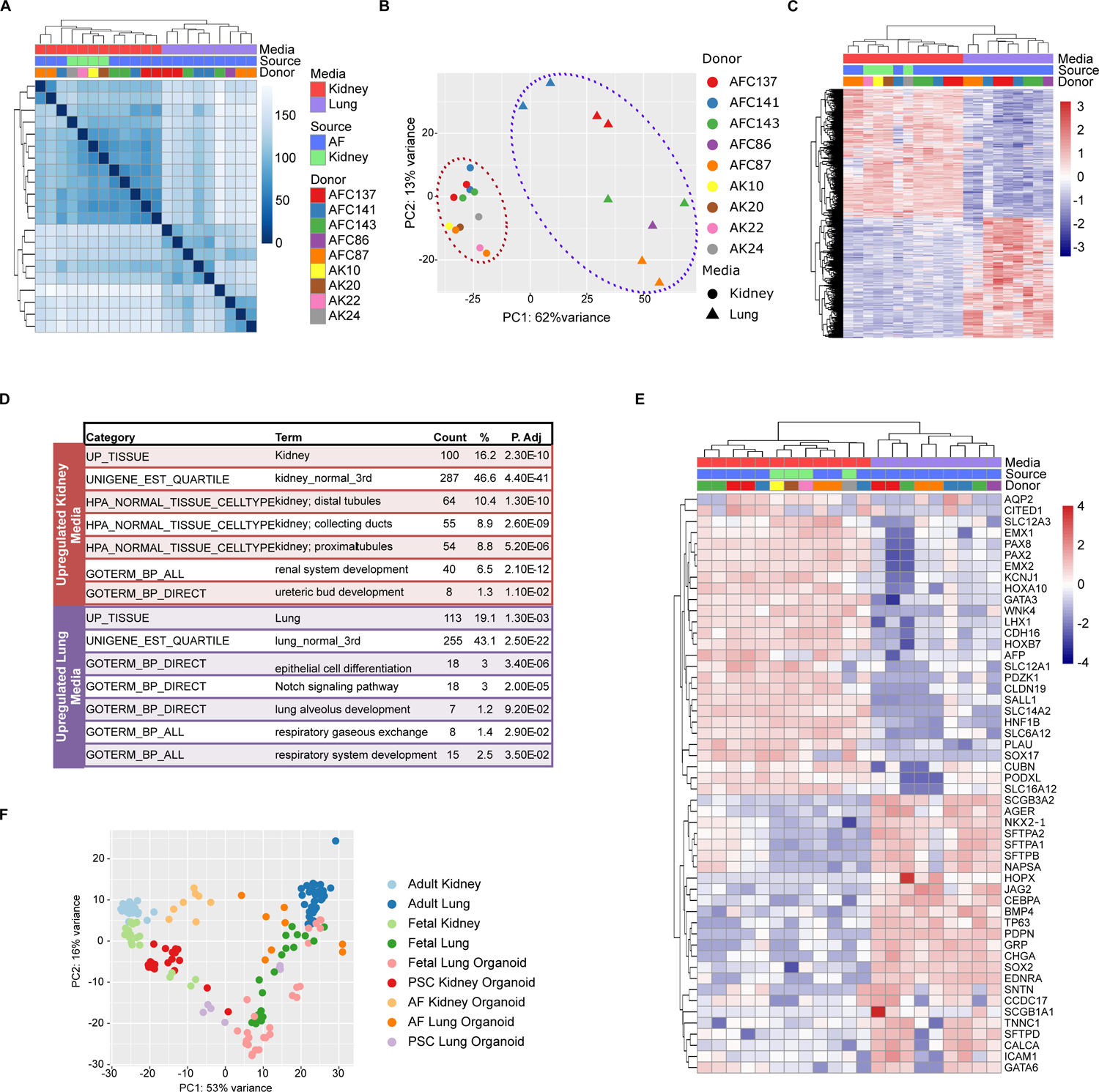
Bulk RNA-seq analysis of AF-derived kidney and lung organoids: **(A)** Heatmap of hierarchical clustering of euclidean distances between indicated samples. **(B)** Principal component analysis (PCA) plot of gene expression in indicated samples. Burgundy and purple dotted ovals indicate Kidney and Lung related organoids, respectively. **(C)** Heatmap of differentially expressed genes between Kidney and Lung related organoids. **(D)** Significantly enriched terms in genes upregulated in Kidney and Lung media. P.adj - p value adjusted by Benjamini-Hochberg FDR correction for multiple testing. **(E)** Heatmap of 52 Kidney and Lung specific genes between Kidney and Lung related organoids. **(F)** PCA plot of gene expression of the genes listed in panel E in the indicated Kidney and Lung related datasets.

We subsequently clustered the samples based on key lung and kidney markers (**Fig. 6E and Sup. Table S3)**. This demonstrated that despite some inter-patient variability, most samples grown in kidney medium, but not samples grown in lung medium, strongly expressed developmental kidney markers (e.g., *PAX2* and *EMX2*), as well as markers of PT (.e.g, *CUBN*), LOH (e.g., *SLC12A1*), DT (e.g., *SLC12A3*) and podocytes (*PODXL*). The opposite was true for the lung medium, which generated organoids expressing markers of lung progenitors (e.g., *SOX2*) and airway (e.g., *TP63*, *SCGB1A1* and *GRP*) and alveolar cells (e.g., *SFTPD* and *ICAM1*).

Next, to better define the transcriptional profile of AF organoids compared to equivalent systems, we used PCA analysis to compare AFKO and AFLO to PSC-derived kidney and lung organoids, fetal kidney and lung tissues and organoids derived from human fetal lungs, based on existing datasets^6,56,93–105^ (**Fig. 6F**). This analysis uncovered that while PSC-derived kidney organoids were more similar to fetal than adult kidneys, as previously described^18^, AFKO seemed to represent an intermediate state between fetal and adult kidneys. Likewise, PSC-derived lung organoids were similar to fetal lungs, whereas AFLO showed an intermediate state between fetal and adult lung. Interestingly, fetal lung organoids (notably comprising several types of systems) showed highly diverse profiles, ranging from fetal-like to adult-like. Altogether, these results show that our defined *in vitro* niches consistently and effectively enrich for either lung or kidney identity, allowing derivation of multi-lineage organ-specific organoids from AF.

## DISCUSSION

Here, we established a platform for prospective derivation of kidney and lung organoids from AF, shown to represent complex multi-lineage replicas of their *in vivo* counterparts. The ability to generate personalized models of fetal organs in real-time has major implications. First, in cases of anomalies detected by prenatal ultrasound, AF-derived organoids could serve as a patient-specific platform to uncover the underlying molecular defect, assess prognosis, and tailor treatments. This could be accomplished during the 2^nd^ trimester, enabling treatment while organogenesis is ongoing, maximizing its effect. Moreover, as amniocentesis is minimally-invasive and already undertaken by many women, such a platform could be readily translated into the clinical realm. This could have a major impact on prenatal medicine, which currently approaches many birth defects as poorly understood and untreatable disorders, managed mostly based on ultrasonographic, rather than molecular findings. Notably, AFKO and AFLO express key genes known to cause congenital anomalies (e.g., *PAX2* and *HNF1B* in AFKO, and *CFTR* and *SFTPB* in AFLO), making them useful tools to study genetic anomalies. Perhaps more importantly, AF organoids offer unprecedented options to study *non-genetic* anomalies, which are currently deemed ‘idiopathic’ in most cases, representing most anomalies^1^. Indeed, these cases can likely be studied only by direct analysis of cells of the affected fetal organ, which until now has been challenging. Moreover, analysis of large AF cohorts, enabled by their lack of ethical constraints and availability, could unravel the molecular underpinnings of development and developmental anomalies across a wide range of genetic backgrounds.

Second, AF organoids could enable deciphering the effect of environmental exposures on development, including infections and drug teratogenicity, for which pre-clinical platforms are lacking^106,107^. Furthermore, it is becoming increasingly clear that environmental exposures during the perinatal period have long-term effects on human health, likely via epigenetic mechanisms, increasing susceptibility to common chronic disorders, a notion termed Developmental Origins of Health and Disease (DOHaD)^4,108^. Although epidemiological studies have uncovered many associations between the prenatal exposome and various health outcomes^109–115^, the mechanistic link is difficult to decipher, due to inaccessibility of fetuses. While researchers have attempted using placenta or cord blood to reflect the fetal epigenome^116,117^, this is likely inaccurate due to the tissue-specific nature of epigenetic signatures^118^. By modeling fetal organs in real-time, AF organoids may allow delineation of these mechanisms, paving the way for uncovering the environmental basis of common diseases, holding promise for improved diagnosis, prognosis and treatment of such diseases.

Third, AF organoids allow studying prematurity complications and elucidating strategies to expedite organ maturation. This is most evident in lung maturation, where steroid treatment has only partial efficacy^90^ and the use of personalized AFLO could determine lung maturity, predict response to steroids, and detect alternative treatments. Nonetheless, other organs represented by AF are also affected by prematurity and may benefit from such a platform, including kidney and colon.

Lastly, AF organoids can be envisioned as a tool for autologous cell therapy using tissue-specific progenitors. Our results, showing AFKO cell integration into the nephrogenic niche as SIX1^+^ NPC may indicate that following longer periods of time, these may give rise to new nephrons. While this requires additional studies, such an approach may be particularly beneficial for severely premature neonates, which have low nephron reserve due to premature termination of nephrogenesis^119,120^. Given their widespread availability, the allogeneic route is also an option in this regard, for instance by generating an AF biobank^121^.

AF organoids represent a new organoid class, adding to adult, fetal and PSC-derived organoids^122^. Similarly to fetal and PSC-derived cells, they model development, as supported by their transcriptome, which is an intermediate between fetal and adult organs. Their advantages over fetal organoids include fewer ethical constraints and availability. Compared to PSC-based approaches, AF organoids are less laborious and reduce the risk for off-target differentiation by relying on lineage-committed progenitors. Moreover, as mentioned above, AF cells are more likely to maintain the epigenetic status of their organ-of-origin, facilitating exploration of DOHaD, as well as idiopathic anomalies. Nevertheless, AF is limited in the tissues that can be modeled, which may also include gastrointestinal organs and skin, making these approaches complementary.

Although AF has long been assumed to contain various fetal cells^123^, there are no high-quality single cell datasets of fresh AF, in part due to low cell viability. Moreover, epithelial cells in AF quickly disappear upon 2D culturing^124^, likely resulting in loss of epithelial progenitors present in AF. Via 3D culture in defined media, we not only preserve these progenitors, but also specifically enrich for either kidney or lung progenitors, which self-organize into tissue-specific organoids. Interestingly, a recent preprint reported derivation of organoids from AF^125^. Although these expressed kidney and lung markers, they were not analyzed at single-cell level, making it hard to determine their exact composition. Notably, rather than using defined *in vitro* niches to prospectively select for the desired progenitors (renal vs. lung) and establish the corresponding organoid type, the employed method relied on retrospective assessment of clones generated from heterogenous AF grown in ‘generic’ media, thus lacking control over which type of organoids will form. Of interest, the same study reported rare formation of organoids expressing intestinal markers, which was seen in 2 samples, only 1 of which was derived from amniocentesis. While this proves the huge potential of AF organoids as a multi-organ platform, it also underscores the necessity of carefully-designed culture conditions so as to control the end-product obtained.

Improper nephrogenesis has major implications. These include: (1) CAKUT, which account for 50% of pediatric end-stage kidney disease^126,127^ and lack specific treatments^128^; (2) Increased risk for kidney disease and hypertension in adulthood, which has been linked to low nephron numbers at birth^129^. Hence, AFKO could have significant translational impact. Several findings regarding AFKO are noteworthy. First, we detected remarkable plasticity of the UB lineage toward tubular phenotypes, corroborating similar plasticity reported in PSC-derived organoids^53,130^, which has yet to be detected *in vivo*. Second, the presence of a stromal fraction is important given the key role of this population in nephrogenesis^43^. Notably, while PSC-derived kidney organoids are rich in stroma^34,35^, adult organoids are strictly epithelial^131^. Third, AFKO harbor urothelial differentiation potential, indicating broad differentiation potential along the UB lineage, including both CD and urothelium, similarly to a recently reported PSC-derived model^54^. This has translational relevance as the majority of CAKUT affect the collecting system. Lastly, we did not detect mature podocytes, yet a podocyte progenitor cluster was noted. This indicates that maturation into differentiated podocytes should be possible, by modifying culture conditions, or by *in vivo* implantation^34,132,133^. Alternatively, integrating vascular elements (e.g., endothelial cells, which are also absent) could promote formation of a glomerular niche with mature podocytes.

AFLO resemble organoids generated from the distal tip of human fetal lungs^6,83^. Intriguingly, we similarly obtain cystic organoids, which later form complex structures, harboring most airway and alveolar cell types. The distal tip initially generates airway cells and gradually switches into an alveolar fate between weeks 17 and 22, the timing of amniocentesis^6,83^. It is thus difficult to determine whether the presence of both airway and alveolar cells reflects *in vitro* plasticity or genuine multipotency of progenitor tip cells. Transcriptomic and TEM analysis revealed that the alveolar cells are immature, expressing both AT1 and AT2 markers, potentially representing a common progenitor. Hence, the organoids could be used to decipher the mechanisms governing lung maturation and identify strategies to enhance it to improve outcomes in prematurity, which accounts for ∼900,000 annual deaths globally^134^. An interesting finding was the presence of a cell type showing common features of both airway and alveolar cells. Although this may reflect *in vitro* cell plasticity, based on recent reports on the presence of such a bipotent cell type in the human adult airways^86,87^, it is possible that these cells actually represent the fetal counterpart of that population. Of note, AFLO lacked mesenchymal cells, which play a key role in lung organogenesis^83^. Improvements in culture media could allow capturing these cells as well, as in AFKO.

In summary, we established a new organoid platform, allowing personalized modeling of the developing fetus during pregnancy. Given its simplicity, lack of ethical constraints, independence from genetic manipulation and reliance on a procedure already undertaken in many patients, it could be readily integrated into clinical practice. Indeed, by affording patient-specific multi-organ AF organoids grown in parallel to the fetus, this work may pave the way to a new paradigm in fetal medicine, replacing conventional AF cultures, and providing a robust way to tailor treatments for fetuses with a suspected anomaly or impending prematurity, two disorders representing a huge clinical burden. Likewise, by providing easy access to organ-specific fetal cells, this work may unlock important insights with implications to diagnosis, prevention and treatment of common chronic diseases that are developmentally programmed. Besides the translational impact, we also expect AF organoids to become a common experimental and pre-clinical tool in a range of disciplines, affording an attractive tool to study the biology of different organs and the deleterious effects of different exposures (e.g., infections or drugs) on development.

## METHODS

### Ethics statements and patient tissue collection

This study was approved by the Sheba Medical Center institutional ethics committee and was performed in accordance with the ethical standards as laid down in the 1964 Declaration of Helsinki and its later amendments (approval numbers 6761-19-SMC, 8993-21-SMC, and 8116-21-SMC). Approved consent was provided by all participants.

### Preparation of RSPO1 conditioned medium

Preparation of RSPO1 conditioned medium was performed from HA-R-Spondin1-Fc 293T Cells according to a distributor’s protocol (R&D systems Catalog# 3710-001-K). The RSPO-1 producing cells were a kind gift from Prof. Benjamin Dekel and Dr. Dana Ishay-Ronen.

### Organoid culture from cells isolated from amniotic fluid

After extraction, amniotic fluid was kept at 4°C before further processing. It was centrifuged at 300 xg at 4°C for 5 minutes, washed 2x with PBS and seeded at a density 50 000 cells/50 μl reduced growth factor BME Type II (Cultrex^TM^, R&D systems) in a 24-well cell culture plate (Corning, Thermo scientific). Alternatively, 12 μl of cell suspension in BME was seeded per well in an 8 chamber slide (Cellvis). After BME polymerization (15 mins at 37°C), cells were incubated in the growth medium (see Supplementary Methods). Medium for AF cells after seeding and organoid cultures after splitting additionally contained ROCK inhibitor Y-27632 (10 μM, Cayman Chemical). The AFKO cultures were split on average every 16 days, while the AFLO cultures were split on average every 21 days. Medium was changed every 3 - 4 days. For splitting, the BME containing organoids was resuspended and sheared with cold base medium, centrifuged at 300 xg at 4°C for 5 minutes, washed once with PBS, and re-seeded in the desired volume of BME.

Cryopreservation was performed by resuspending the BME containing organoids with cold base medium, washing once with PBS, and resuspending in a freezing solution composed of 90% FBS (Gibco), 10% DMSO (Sigma). The organoids were frozen using a cryo cooler at −80°C and stored at −80°C.

### Organoid culture from cells isolated from adult kidney

Human nephrectomy tissue specimens were used to establish tubuloids as described previously^92^. In brief, the tissue was dissected and minced; the fragments were digested with 1 mg/ml of Collagenase (C9407, Sigma) in combination with 100 µg/ml Liberase TL (540102000, Sigma) in DMEM/F12 medium (Sartorius) for 45 min at 37°C with shaking at 160 RPM using a thermal shaker. The digested fragments were passed through a 100 μm cell strainer (Greiner). The cells were then washed twice with PBS, resuspended in BME (50,000 cells/50 μl BME drop) and seeded in a 24-well cell culture plate (50 μl BME/well). BME was incubated for 15 minutes at 37°C to solidify, and kidney medium was added.

### Immunostaining and imaging

Whole-mount staining of organoids was performed in an 8 chamber slide (Cellvis), each well containing 12 μl of BME with organoids. Before fixation, the organoids were washed once with PBS for 5 mins, fixed using 4% paraformaldehyde (PFA) for 1 hour at room temperature.

Permeabilization was performed using 0.3% Triton-X in PBS for 30 mins, blocking for 1 hour with 3% BSA dissolved in 0.3% Triton-X in PBS. After blocking, primary antibodies diluted in the blocking solution were incubated overnight at 4°C, shaking 60 RPM. The list of antibodies can be found in Supplementary methods. After primary antibody incubation, organoids were washed 3x with 0.3% Triton-X in PBS, 1 hour per wash. Secondary antibodies diluted in the blocking solution were then incubated overnight at 4°C, shaking 60 RPM. Phalloidin-iFluor 647 Reagent (ab176759, Abcam) was incubated alongside the secondary antibodies. After secondary antibody incubation, organoids were washed 3x with 0.3% Triton-X in PBS, 1 hour per wash, followed by 1 µg/ml DAPI (Sigma) incubation for 10 minutes at room temperature. After a final 5 minute wash with 0.3% Triton-X in PBS, the organoids were incubated in PBS for imaging with a Zeiss LSM 700 microscope. The images were processed using Zen 2.5 (blue addition) software.

For immunofluorescence staining of paraffin sections, the organoids were fixed in 4% PFA, embedded in bio agar (05-9803S, Bio-Optica) and then fixed again in 4% PFA overnight. Tissues were also fixed in 4% PFA overnight. After fixation, samples were dehydrated and embedded in paraffin. Deparaffinization and antigen retrieval was performed using Omniprep Solution pH 9 (Zytomed Systems) according to manufacturer’s instructions on 5 µm sections. Slides were then incubated for 20 minutes in CAS-Block (Invitrogen). Afterwards, the slides were incubated with primary antibodies diluted in Antibody Diluent (ThermoFisher Scientific) for 1 hour at room temperature, washed twice with 0.1% Tween in PBS, incubated with secondary antibodies for 1 hour at room temperature and washed twice with PBS. Lastly slides were closed with DAPI FLUOROMOUNT G (Electron Microscopy Sciences) and subjected to microscopy. Olympus IX83 widefield system equipped with a DP80 camera and driven by cellSens version 4.2 was used for image acquisition. OLYMPUS OlyVIA 3.4.1 software was used for image processing.

### Transmission Electron microscopy (TEM)

Organoids were fixed in Karnovsky-fixative (2% glutaraldehyde, 3% paraformaldehyde, in 0.1M cacodylate-buffer pH 7.4), for 2.5 hours at room temperature, washed 4 times in 0.1M cacodylate-buffer pH 7.4. The tissue was embedded in 3.4% agarose post-fixation in 1% OsO4, contain 0.5% K2Cr2O7, 0.5% K4[Fe(CN)6]3H2O, in 0.1M cacodylate-buffer for 1 hour at room temperature and washed in cacodylate buffer 0.1 M and three times in DDW. Then stained with uranyl-acetate in DDW (2% for 1 hour) in the dark, washed twice in DDW. After a series of dehydrating steps in rising EtOH-concentrations (50%, 70%, 96% each twice 5 minutes, 100% three times 10 minutes, followed by 2 times propylene oxide, 5 minutes). Epon (“hard”) was introduced gradually into the tissue in rising concentrations (30%, 50%, 70%) 2 hours each, 50% overnight and again 100% three times for two hours). The blocks were incubated at 70°C for 72 hours. The blocks were then cut to 70-100 nm ultra-thin sections with ultra microtome Leica EM UC7 and stained with lead citrate for 4 min. Sections were imaged with a JEOL 1400 With Gatan CCD camera 2kx2k.

### Reverse-transcriptase quantitative PCR

Total RNA was purified from organoids using Tri-Reagent (Sigma) or Directzol RNA Microprep kit (Zymo research). 500 nanograms of RNA was used as a template to synthesize cDNA using the Verso cDNA Synthesis Kit (Thermo Scientific). RT–qPCR was performed in triplicate with cDNA (1:10 dilution), 250 nM primers and Fast SYBR™ Green Master Mix (Applied Biosystems) using the StepOnePlus Real-time PCR System (Applied Biosystems). The data were analyzed using StepOne Software v2.3. *ACTB* was used as the endogenous control. For the list of primers, see Supplementary Table 4.

### MARS-seq RNA sequencing and analysis

Total RNA was extracted from organoids using either Tri-reagent (Sigma) or Directzol RNA Microprep kit (Zymo research) following the manufacturer’s instructions. 3’ RNA-seq (Bulk MARS-seq^91,135^) libraries were prepared and sequenced on a Novaseq 6000 (Illumina) at the Weizmann Crown Institute for Genomics. Poly-A/T stretches and Illumina adapters were trimmed from the reads using cutadapt^136^; resulting reads shorter than 30bp were discarded. The remaining reads were mapped onto 3’ UTR regions (1000 bases) of the H. sapiens, GRCh38_p13 genome according to ensembl annotations v 106, using STAR^137^, with EndToEnd option and outFilterMismatchNoverLmax was set to 0.05. Deduplication was carried out by flagging all reads that were mapped to the same gene and had the same UMI. Counts for each gene were quantified using htseq-count^138^, using the gtf above. Only uniquely mapped reads were used to determine the number of reads that map to each gene (intersection-strict mode). UMI counts were corrected for saturation by considering the expected number of unique elements when sampling without replacement. Gene count matrices were analyzed with DEseq2 (Build 1.38.3) in R (version 4.2.2). Genes with low counts across samples were filtered out and differentially expressed genes between organoids grown in kidney media (both tubuloids from adult kidney donors and AFKOs) and organoids grown in lung media (AFLOs - limited to lower passage samples) were uncovered with the results() function, alpha = 0.05, lfcThreshold = 0.58.

Functionally enriched terms were found within the differentially expressed gene lists using DAVID https://david.ncifcrf.gov/. The counts underwent RLOG transformation and similarity between samples was measured by hierarchical clustering of euclidean distances between samples’ RLOG transformed counts, as well as principal component analysis. Heatmaps were produced with pheatmap function, with differentially expressed gene heatmaps deriving from the RLOG transformed counts, normalized by row. In order to compare MARS-seq data to previously published RNAseq datasets of adult, fetal and pluripotent stem cell and organ derived organoids of lung and kidney, the MARS-seq counts were normalized to TPM, and TPM data was downloaded from the GEO repository (Adult Kidney - GSE93480^93^, GSE102101^94^, GSE106548^95^, GSE213324; Fetal Kidney - GSE75949, GSE100859^56^; PSC Kidney Organoid - GSE145085^96^, GSE164648, GSE182350; Adult Lung - GSE86958^97^, GSE92592^98^, GSE102511^99^, GSE133206^100^; Fetal Lung - GSE83888^101^, GSE95860^6^, GSE121238^102^; PSC Lung Organoid - GSE148697^104^, GSE151796^103^; Fetal Lung Organoid - GSE211308^105^, GSE83888^101^, GSE95860^6^). TPM values of Adult, Fetal and AF organoids for Kidney and Lung samples on specified genes were directly compared by student’s t-test using rstatix package (version 0.7.2), with p-values adjusted by Holm method and graphs created with ggpubr package (version 0.6.0) in R. All datasets were compared with principal component analysis focusing on RLOG transformed TPM values of lung and kidney specific marker genes.

### Single cell RNA sequencing

The processing of organoids into single cells is described in the Fetal kidney explant injection section. After TryPLE digestion, the cell suspension was passed through a 40 μm strainer (Greiner). Subsequently cells were washed twice with 0.04% BSA in PBS and kept on ice until further processing. Single cells were captured and barcoded on a Chromium Controller (10X Genomics). Subsequently, RNA from the barcoded cells was reverse-transcribed and sequencing libraries were prepared using Chromium Single Cell Reagent Kit v3.1 Chemistry Dual Index (10X Genomics) according to the manufacturer’s instructions. Libraries were sequenced on a Novaseq 6000 machine at the Weizmann institute.

### Analysis of scRNA-seq data

CellRanger pipeline^139^ (v6.0.1, 10x genomics) with default parameters was used for demultiplexing, alignment (hg38 reference genome, 2020-A version, downloaded from 10X website), filtering, barcode counting, and UMI counting. The Seurat R package^140^ (v4.0.4) was used for downstream analysis and visualization. Gene-cell matrices were filtered to remove cells with 4MADs above the median of mitochondrial genes and with less than 250 genes and 500 UMIs. In addition, genes detected in fewer than 5 cells were excluded from the analysis. After implementing these quality control measures, a total of 3,675 AFKO123 cells, 5,580 AFKO133 cells, 7,349 AFLO131 cells, and 18,752 AFLO171 were retained for further analysis.

The expression data was normalized using Seurat’s NormalizeData function, which normalizes the feature expression measurements for each cell by the total expression, multiplies this by a scale factor (10,000), and then log-transforms the results. The top 2,000 highly variable genes were identified using Seurat’s FindVariableFeatures function with the ‘vst’ method. The four Kidney and Lung samples were then integrated using Seurat’s integration functions^141^. Briefly, FindIntegrationAnchors function was used to identify anchors between the two samples and an integrated assay was created using the IntegrateData function. Further analysis was done on the integrated Kidney and Lung samples. Each cell was assigned with a cell-cycle score using the CellCycleScoring function and the G2/M and S phase markers lists from the Seurat package. Potential sources of unspecific variation in the data were removed by regressing out the mitochondrial gene proportion and UMI count using linear models and finally by scaling and centering the residuals as implemented in the function “ScaleData” of the Seurat package. Principal component analysis (PCA) was performed. 18 PCs for both the Kidney and Lung datasets, were used for clustering and data reduction. Cell clusters were generated using Seurat’s unsupervised graph-based clustering functions “FindNeighbors” and “FindClusters” (resolution = 0.5). UMAP was generated using the RunUMAP on the projected principal component (PC) space. Seurat’s functions FeaturePlot and DimPlot were used for visualization. Seurtat’s DotPlot function was used to generate dot plots to visualize gene expression for each cluster. Plots were further formatted using custom R scripts with the packages ggplot2^142^ and patchwork^143^ Heatmaps were produced with Seurat’s function DoHeatmap or the R package pheatmap^144^.

Marker genes for each cluster were identified by performing differential expression between a distinct cell cluster and the cells of the other clusters with the non-parametric Wilcoxon rank sum test (Seurat’s FindAllMarkers function). Cell types were assigned manually based on the expression of classic marker genes. Cells from cluster 8 from integrated Kidney analysis were assigned as Lung cluster and were removed from downstream analysis.

Organ of origin was determined by the expression of organ specific genes as listed in Supplementary Table 1. A gene was considered expressed if it had an expression of >0.3 in the particular cluster. Every expressed gene was given a value 1, while non-expressed genes were given value 0. The normalized expression is the ratio of organ-specific gene values to the sum of all the values in the particular cluster.

Two subsets of the integrated Kidney samples (subset1 - clusters 0,4,5,7,8,11,12 and subset2 – clusters 1,2,3,6,9) and the integrated Lung sample were re-analyzed for trajectory analysis using monocle3 R package 4. The following functions were applied for preprocess, batch correction ^145^ and data reduction – preprocess_cds with 50 dimensions, align_cds with samples as alignment_group and reduce_dimension with ‘Aligned’ as preprocess_method. Cells were clustered using the cluster_cells function (res 0.002) and the trajectory graph was learned using the learn_graph function. Pseudotime was calculated using the order_cells function. CellPhoneDB^146^ was used to find interactions between clusters. Results were plotted using the R package ktplots^147^.

## Data availability

NGS data will be made available upon publication at the NCBI GEO repository.

## ACKNOWLEDGEMENT

We thank Dr. Ronnie Blecher from the Crown Genomics institute of the Nancy and Stephen Grand Israel National Center for Personalized Medicine, Weizmann Institute of Science for assistance in preparation and sequencing of scRNA-seq libraries, Dr. Avital Sarusi-Portuguez from the Mantoux Bioinformatics institute of the Nancy and Stephen Grand Israel National Center for Personalized Medicine, Weizmann Institute of Science for assistance in scRNA-seq data analysis, Helena Sabanay from the Bar Ilan institute of Nanotechnology and Advanced Materials for assistance with transmission electron microscopy. We thank Dr. Dana Ishay-Ronen for the gift of R-spondin expressing cells and breast organoids, and Prof. Benjamin Dekel’s lab for assistance with microscopy experiments. O.P. is supported by the Sheba Medical Center Physician-Researcher Program, Israel Science Fund (grant no. 2224/21), U.S.-Israel Binational Science Foundation (grant no. 2021164), Israel Ministry of Science and Technology (grant no. 0004571), Estates Committee of Israel Ministry of Justice (grant no. 20230670), Alrov Fund, and the Suzanne Eichinger-Henke Fund (Faculty of Medicine, Tel Aviv University). R.L-K is partially supported by The Nehemia Rubin Excellence in Biomedical Research, TELEM Program, Sheba Medical Center, Tel Hashomer, Israel. B.W. is supported by the Israel Ministry of Science and Technology (grant no. 0004571), Estates Committee of Israel Ministry of Justice (grant no. 20230670). P.B is supported by the Recanati Grant (Faculty of Medicine, Tel-Aviv University, Israel).

## Author Contributions

O.B., B.W., R.L.-K., and O.P. conceptualized the overall project. O.B., R.L.-K., and G.R. conducted AFKO cell culture experiments, immunostainings and microscopy. O.B. conducted qRT-PCR and functional assays on AF derived organoids. O.B., and G.R. performed fetal kidney explant injections. T.J. assisted with adult kidney tubuloid extraction. O.B., R.L.-K., and O.P. performed formal and computational analysis of data. H.A., H.S., A.S., L.B., N.P., T.A., D.S., D.D.F., I.B., Z.A.D, and B.W. helped with data acquisition. O.P., R.L.-K., B.W., and P.B. contributed to funding. O.B., R.L-K., B.W., P.B., and O.P. drafted the manuscript, with all authors contributing to manuscript proofing and revision. R.L.-K. and O.P. supervised the work.

